# Acoustofluidic patterning for improved microtissue histology

**DOI:** 10.1101/2025.02.09.637288

**Authors:** Dhananjay V. Deshmukh, Emilie Vuille-dit-Bille, Saumitra Joshi, Nicola Gerber, Christine Fux, Annina Eichenberger, Epifania Bono, Markus Rimann, Sarah Heub, Daniel Migliozzi, Stephanie Boder-Pasche, Gilles Weder, Jurg Dual, Mark W. Tibbitt

**Affiliations:** Macromolecular Engineering Laboratory, Department of Mechanical and Process Engineering, ETH Zurich, Zurich, Switzerland; Institute for Mechanical Systems, Department of Mechanical and Process Engineering, ETH Zurich, Zurich, Switzerland; CSEM SA, Neuchâtel, Switzerland; Institute of Mechanical Engineering, EPFL, Lausanne, Switzerland; 3D Tissues and Biofabrication, Institute of Chemistry and Biotechnology, Zurich University of Applied Sciences (ZHAW), Waedenswill, Switzerland; Competence Center TEDD, Institute of Chemistry and Biotechnology, Zurich University of Applied Sciences (ZHAW), Waedenswil, Switzerland

## Abstract

Microtissues (such as organoids or spheroids) are useful to model healthy or diseased human tissues and organs, with application in drug testing, tissue engineering, and personalized medicine. To characterize their morphology and phenotype, microtissues are often monitored using histology and high-resolution imaging. While this technique is well-established and standard for macroscopic tissue sections, it remains challenging to execute and has low throughput in the context of these sub-millimeter structures, mainly due to the fact that their size makes them difficult to locate within the embedding medium. To enhance the efficiency of microtissue histology, we developed an acoustofluidic device that arranges microtissues in a coplanar arrangement within HistoGel, a widely used embedding medium. Pre-patterning microtissues within HistoGel streamlines downstream processing by positioning the centers of mass of the microtissues at a pre-defined plan in the sample, simplifying sectioning, labeling, and imaging. We validated the method using differently sized HepG2 microtissues and tumor microtissues from osteosarcoma cells (Saos-2, MG-63, HOS), demonstrating that this method can be applied to different microtissue shapes, sizes, and types, without modifying established post-processing techniques. Our approach substantially enhances the microtissue histology workflow and broadens possibilities for microtissue analysis.

## Introduction

Multicellular assemblies can self-organize into three-dimensional (3D) structures that recapitulate native tissues and organs. Intestinal organoids exhibit crypt-like structures^1^; cerebral organoids capture brain development^2^; liver organoids produce digestive enzymes^3^; and retinal organoids develop functional photoreceptors^4^. These microtissues (an umbrella term for spheroids, organoids, tumoroids, assembloids, and other multicellular assemblies) have transformed biomedical research by providing physiologically relevant models at the micron-scale to study tissue function, monitor disease progression, and screen potential therapeutics, enabling a smoother transition of laboratory discoveries into medical applications^5,6^.

The uses of microtissues in disease modeling and drug testing are particularly impactful, as they offer critical advantages over traditional in vitro methods^7^. This is primarily based on their improved phenotypic and biological relevance as compared with 2D monocultures. For instance, liver organoids were used to model steatohepatitis, a liver disease characterized by steatosis, inflammation, and fibrosis^8^. The liver organoids reproduced disease progression, providing insight into its pathology, and were used to screen potential drugs to treat steatohepatitis. In parallel, patient-derived microtissues enable personalized medicine by recapitulating disease heterogeneity in a dish, such as for rhabdoid tumors in children^9^ and osteosarcoma, a rare but complex and heterogenous cancer with high chemotherapy resistance^10–12^. Mimicking and then studying the molecular and cellular complexity of such diseases is particularly relevant to enhance the applicability of drug screening assays^10,13^. Given the limited availability of patient-derived cells, optimized workflows for analyzing microtissues are critical to minimize sample loss and maximize clinical insight.

Despite substantial advances in microtissue production, including methods like hanging drop culture^14^, low-adhesion substrates^15^, and micro-molded surfaces^16^, challenges remain in the monitoring and analysis of microtissues. Optical imaging techniques, such as light-sheet microscopy and sample clearing, have dramatically improved visualization but remain low throughput and incompatible with standard histological protocols^17^. Limited tools to manipulate and monitor microtissues downstream remain as roadblocks in their translation and clinical utility^18^. Specifically, while histology remains the primary method for analyzing microtissues in the clinical setting, the process is inefficient and cumbersome, especially for patient-derived microtissue samples.

Microtissue histology necessitates that the samples be suspended in an embedding medium, such as HistoGel (an agarose-based gel commonly used for microtissue processing) prior to being sectioned, stained, and imaged to characterize their structure and phenotype^19^. The embedding process introduces inherent challenges due to the small size of the microtissues and the random distribution within embedding media. The stochastic placement of microtissues within the gel means that a large fraction of the samples needs to be sectioned and imaged to locate and analyze the samples, increasing the time needed for and cost associated with analysis as well as complicating traceability across sections^20^.

Without pre-organization of microtissues in the embedding gel, the histological process is time and material consuming and can lead to loss of precious samples and the data associated with those samples.

Some technologies have been developed to improve microtissue histology processing. Passive approaches, such as embedding microtissues in micropatterned gels, have improved sectioning efficiency and reduced sample loss^20,21^. However, these methods are less effective for heterogeneous microtissues, which vary in size or shape, and make it challenging to pre-define a plane for sectioning to capture all samples. Active positioning methods, such as acoustofluidics, would offer a more versatile solution. Acoustofluidics uses contactless, label-free acoustic forces to manipulate cells or microtissues in liquid media safely and precisely^22^. To enable acoustic manipulation, a standing pressure wave can be generated within a liquid medium. When cells or microtissues are suspended in this acoustic field, they move towards the pressure nodes. Unlike passive methods, which rely on settling microtissues to the bottom of preformed gels, acoustofluidics enables precise alignment of microtissue centers of mass to a pre-defined plane, mitigating misalignment caused by size heterogeneity or steps in sample alignment. Acoustofluidics has already been used to produced microtissues^23^ and to orient non-spherical structures^24^ and spheroids^25^. Acoustic forces scale with object size^26^, making this approach particularly effective for positioning larger structures like microtissues.

Building on the advantages of acoustofluidics, we developed a device to pre-pattern microtissues within HistoGel for improved histological analysis. By integrating temperature control, we manipulated the viscosity of HistoGel for acoustofluidic alignment in its liquid state and solidified the gel to fix the positions of microtissues in pre-defined, co-planar arrangements. The patterned samples were removed and embedded into a thicker slab of HistoGel for multiplexing and downstream processing, sectioning, and imaging, using previously established standard protocols. This process reduces the number of sections needed for analysis, minimizes sample loss, and allows multi-sample arrangements for parallel processing of patient-derived microtissues. Validation using liver and osteosarcoma microtissues demonstrated the robustness and versatility of our approach across diverse microtissue sizes and shapes. This technique has the potential to streamline histology workflows, enhancing the efficiency and utility of microtissues in biomedical research.

## Results and discussions

### Acoustofluidic device enables microtissue levitation

Despite its critical role in assessing microtissue biology, conventional microtissue histology is inefficient due to the random distribution of the small biological samples within the embedding medium (**Figure 1a**). This results in the need to section the whole sample even though the biological specimens are only present in a limited fraction of the total volume, making downstream analysis time-consuming and costly^27^. To address this challenge, we leveraged acoustofluidics to organize microtissues so that they are coplanar at a defined height in the final sample, enabling the user to locate the microtissues easily and increase the efficiency of analysis as all microtissues are captured in a small number of slices (**Figure 1b**). Further, this process was designed so that the contactless patterning using acoustofluidics would proceed within standard embedding medium and be compatible with downstream histological processing and imaging.

**Figure 1.**
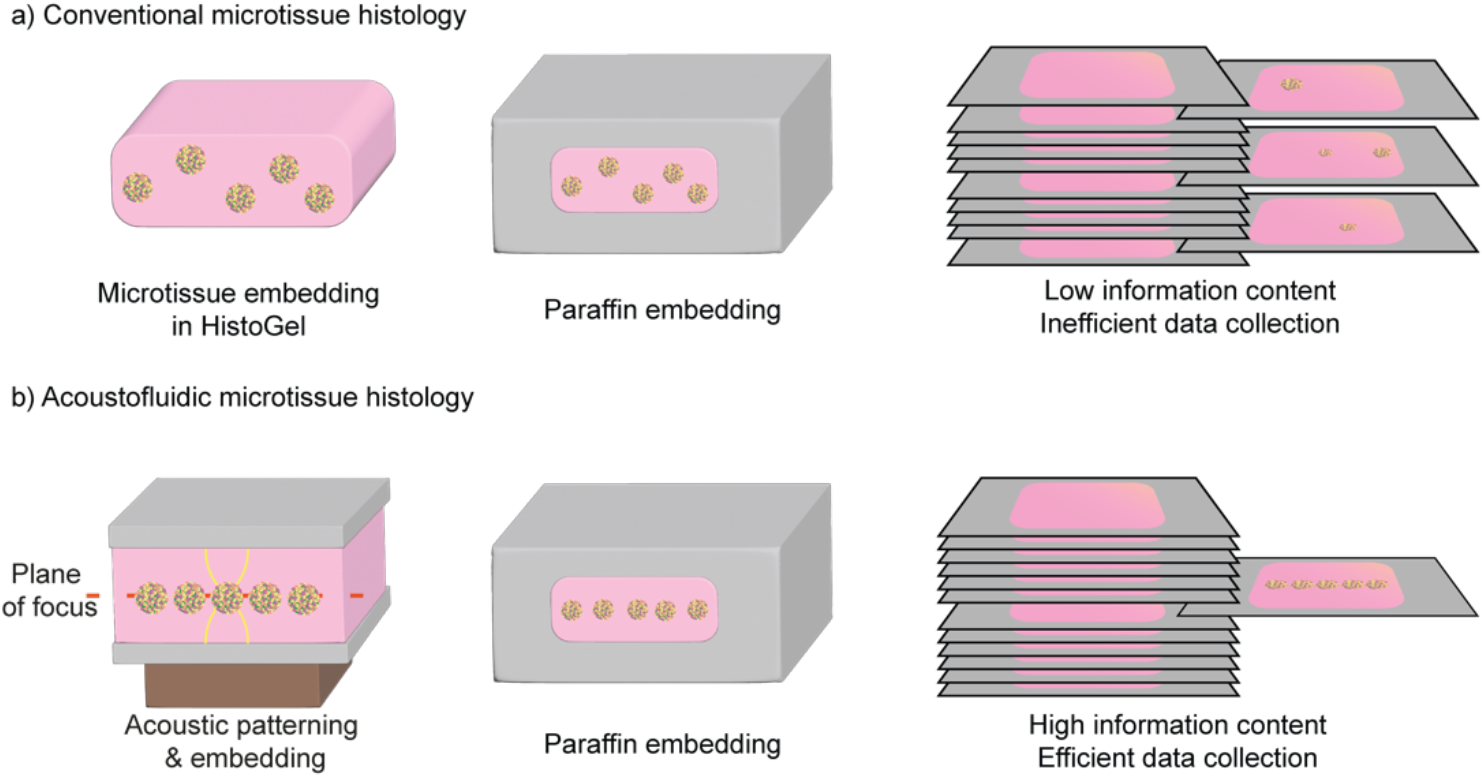
Increased microtissue histology efficiency via coplanar patterning using acoustofluidics. **a)** In conventional microtissue histology, microtissues are randomly distributed in the sample which makes their analysis inefficient. **b)** Using acoustofluidics microtissues can be patterned such that they are coplanar, enabling fast and efficient data collection.

To enable contactless patterning of microtissues, we designed an acoustofluidic device consisting of a fluidic cavity in an aluminum plate with a piezoelectric transducer attached on one side (**Figure 2a,b**). The piezoelectric transducer was actuated using an electric signal with controllable frequency and voltage, which allows control over spatial profile and the intensity of the pressure field inside the acoustic cavity. Given the importance of precisely positioning the microtissues to a single plane in our application, an acoustic cavity was designed to ensure precise control over the pressure fields. We selected an acoustic cavity with a rectangular cross-section (W x H: 1.1 mm x 1.5 mm) and length of 30 mm, with a piezoelectric transducer mounted on the side to levitate the microtissues along the central plane of the cavity. The rectangular cross-section was chosen over the square cross-section as the asymmetric cross-section resulted in more reliable and reproducible acoustofluidic patterning (**Figure S1**).

**Figure 2.**
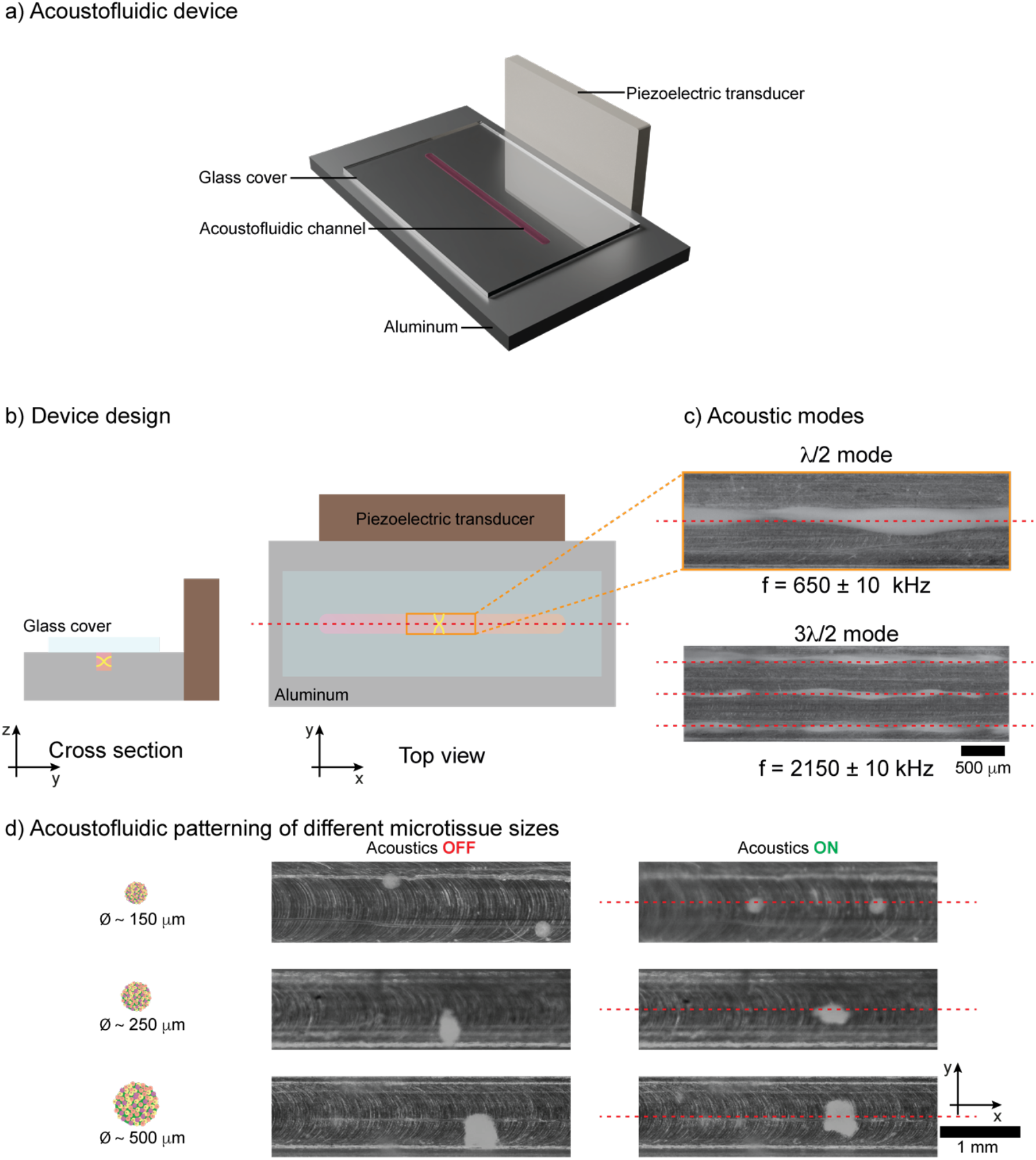
Acoustofluidic device for microtissue manipulation. **a)** Device design. **b)** Vibrations from the piezoelectric transducer produces a standing wave in y-direction, resulting a pressure node in the center of the channel. **c)** Acoustic characterization of the device using yeast, the first and third modes were observed at 650 kHz and 2150 kHz respectively. **d)** Different sizes of HepG2 microtissues can be patterned with the device. Dashed red lines (in b, c, and d) show expected pressure nodes.

The resonant frequency of the device was identified via a frequency sweep with tracking particles (yeast) to identify the resonant frequency of the device. The resonant frequencies for the 1^st^ (λ/2) and 3^rd^ (3λ/2) modes were identified as 650 kHz and 2150 kHz, respectively (**Figure 2c**). To account for changes in resonant frequency caused by experimental variations in temperature or device clamping, we applied a frequency-modulated signal across the piezoelectric transducer in the subsequent patterning experiments. The modulation used a sine wave with a frequency deviation (Δf) of 20 kHz and a modulation frequency of 100 Hz.

Once the resonant frequencies and the operating parameters were identified, we patterned fixed HepG2 microtissues in PBS using the acoustofluidic device. We confirmed that the same acoustic parameters were capable of patterning microtissues **(Figure S1, S2)**. The acoustic field was applied just before loading the device with microtissues. A glass slide was then used to cover the device and enclose the acoustic cavity (**Figure S2**). While all the above experiments were done with similarly sized microtissues, we also tested our method for differently sized microtissues (Ø ∼ 150, 250, and 500 μm). We found that patterning of the microtissues can also be done with different sizes of HepG2 microtissues using the same protocol (**Figure 2d**). In these experiments, we further observed that non-spherical microtissues were successfully patterned and that their orientation changed in response to the acoustic field, orienting perpendicular to the pressure waves.

### Acoustofluidic patterning of microtissues within HistoGel

To integrate this method into existing microtissue histology workflows, it is essential to be able to pattern microtissues in standard embedding media, such as HistoGel an agar-based embedding medium commonly used for microtissue histology. HistoGel becomes a liquid when heated above 60 °C, which allows for acoustofluidic levitation of the microtissues. When cooled down, it turns back into a solid gel embedding the microtissues and holding their patterned position. HistoGel displays temperature hysteresis, such that after heating it will remain a liquid until about 40 °C, however once solidified, HistoGel does not liquify again until it reaches ∼60 °C. To use our device with HistoGel, we integrated an arduino-based temperature control system in the device that enabled us to monitor and control device temperature (**Figure S3a**). The device could be heated to 45 °C in ∼5 min and cooled (using a cold metal block) to 10 °C in ∼4 min (**Figure S3b**). The elevated temperature (45 °C) was sufficient to maintain HistoGel in a liquid state—after being heated to 60 °C outside of the device—until microtissue patterning was complete.

Compared with our characterization experiments in aqueous media, there are two important differences when microtissues were patterned within HistoGel. First, the change in viscosity affected the Stokes drag experienced by the microtissues. Second, due to the change in the medium the speed of sound differed, meaning that the resonant frequencies of the fluid cavity may shift slightly. For the patterning experiments, we used a diluted HistoGel (65 vol% in PBS), to limit changes in viscosity and speed of sound while still allowing for embedding via solidification and downstream processing. The viscosity of our HistoGel solution (∼85 mPa.s at 45 °C; **Figure S4**) was orders of magnitude higher than that of water. To counter the increase in the drag forces the voltage applied across the piezoelectric transducer was increased (from 20 Vpp for water to 30 Vpp for HistoGel). The speed of sound in our HistoGel solution was 1548 m s^-1^ at 60 °C and 1535 m s^-1^ at 45 °C, which was higher than the speed of sound in water (1498 m s^-1^) but the difference was subtle and did not require altering the applied frequencies since frequency modulation in our signal already accounted for small changes in the resonant frequencies due to the changes in the speed of sound (**Figure S5**).

Microtissues were patterned to the center of the acoustofluidic device. A patterning efficiency of 79.8 ± 7.63 % was obtained over 14 experiments involving over 800 HepG2 microtissues. In comparison, when acoustofluidics was not applied, the patterning efficiency dropped substantially to 11.51 ± 5.99% (**Figure 3a**), consistent with random placement in the volume. To further validate our techniques, we patterned three types of osteosarcoma microtissues, HOS, Saos-2, and MG-63 (**Figure 3b**), which differ in size, shape, and compactness. Remarkably, the same operating parameters and protocol were effective across all microtissue types, demonstrating the versatility of our approach for microtissues with different sizes and properties.

**Figure 3.**
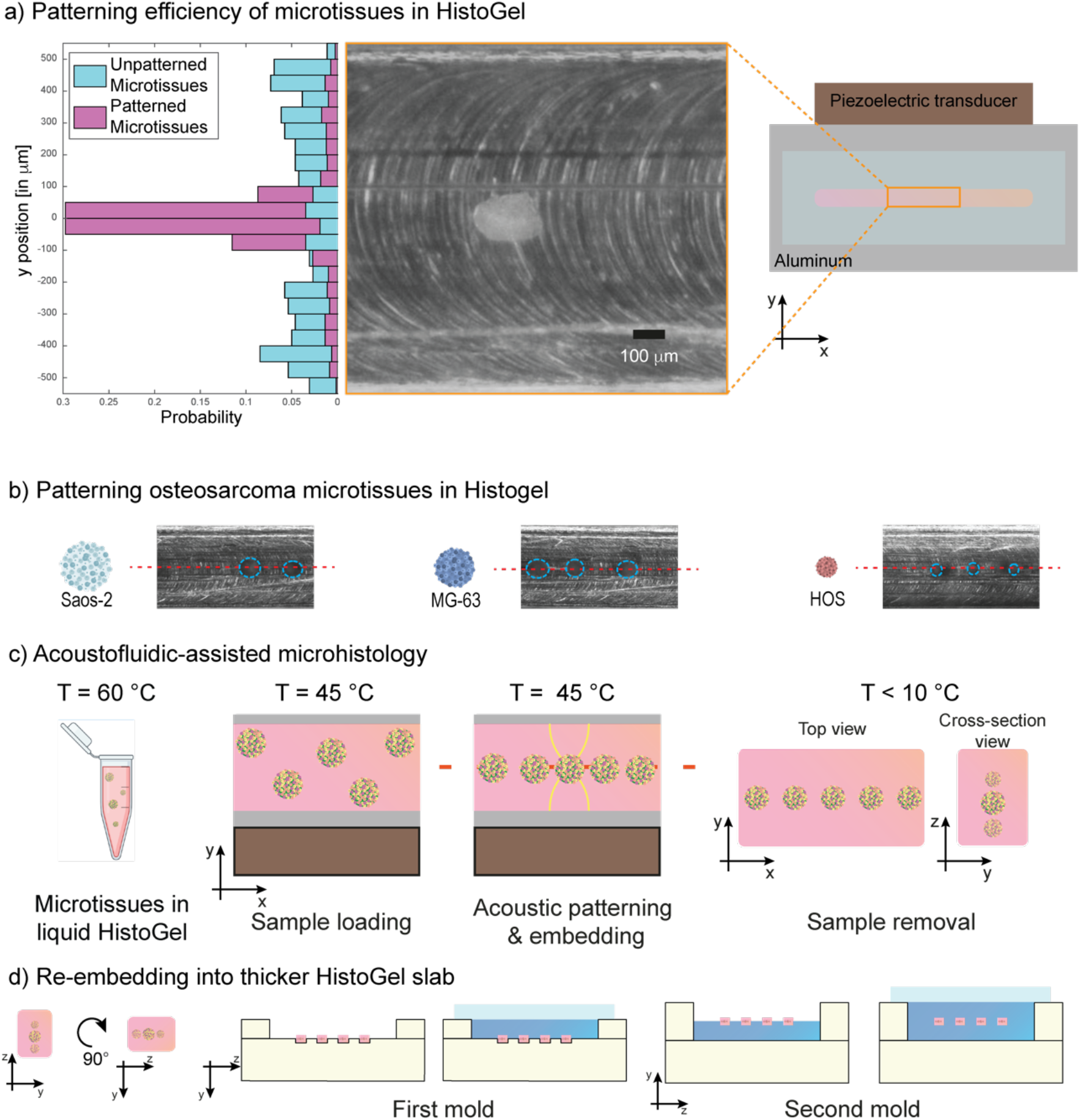
Acoustofluidic patterning in HistoGel and post-processing. **a)** Patterning efficiency of microtissues in HistoGel comparing acoustically patterned (pink) with unpatterned and random positioning (blue). **b)** Osteosarcoma microtissues (Saos-2, MG-63, and HOS) patterned within HistoGel. Microtissues are outlined with a cyan dashed circle, while the center of the channel is indicated with a red dashed line. **c)** Process steps for acoustic patterning of microtissues within HistoGel. **d)** Re-embedding the acoustofluidically patterned samples (pink) into a thicker slab of HistoGel (blue). The surrounding HistoGel was dyed blue to distinguish it from the samples. Glass slide (in light blue) was used to cover the HistoGel slab during molding to ensure a flat top and bottom surface.

### Acoustofluidic patterning enables efficient sectioning of microtissue samples for histological analysis

Having established that acoustofluidics can be used to pattern microtissues in a coplanar fashion within HistoGel, we applied our method for multiplexing in standard histology cassettes (**Figure 3c,d**). The patterned microtissue samples were further embedded within a dyed HistoGel outer layer in a sequential process to clearly distinguish between the sample of interest (within undyed HistoGel) and the surrounding HistoGel (dyed). We combined four distinct acoustically patterned samples such that the centers of mass of all microtissues were in the same plane within the final multiplexed sample. Each of the four samples were placed 1.5 mm from each other and were embedded in 6 mm thick HistoGel. These thicker embedded constructs were easier to handle and maintained better geometric integrity during the post-processing (**Figure 4a**). Overall, this method streamlined post-processing and improved the reproducibility of comparative analyses across different microtissue samples or treatment conditions. Further, multiplexing samples within each construct simultaneously reduced time and resource consumption, ultimately enhancing the throughput and reliability of the results.

**Figure 4.**
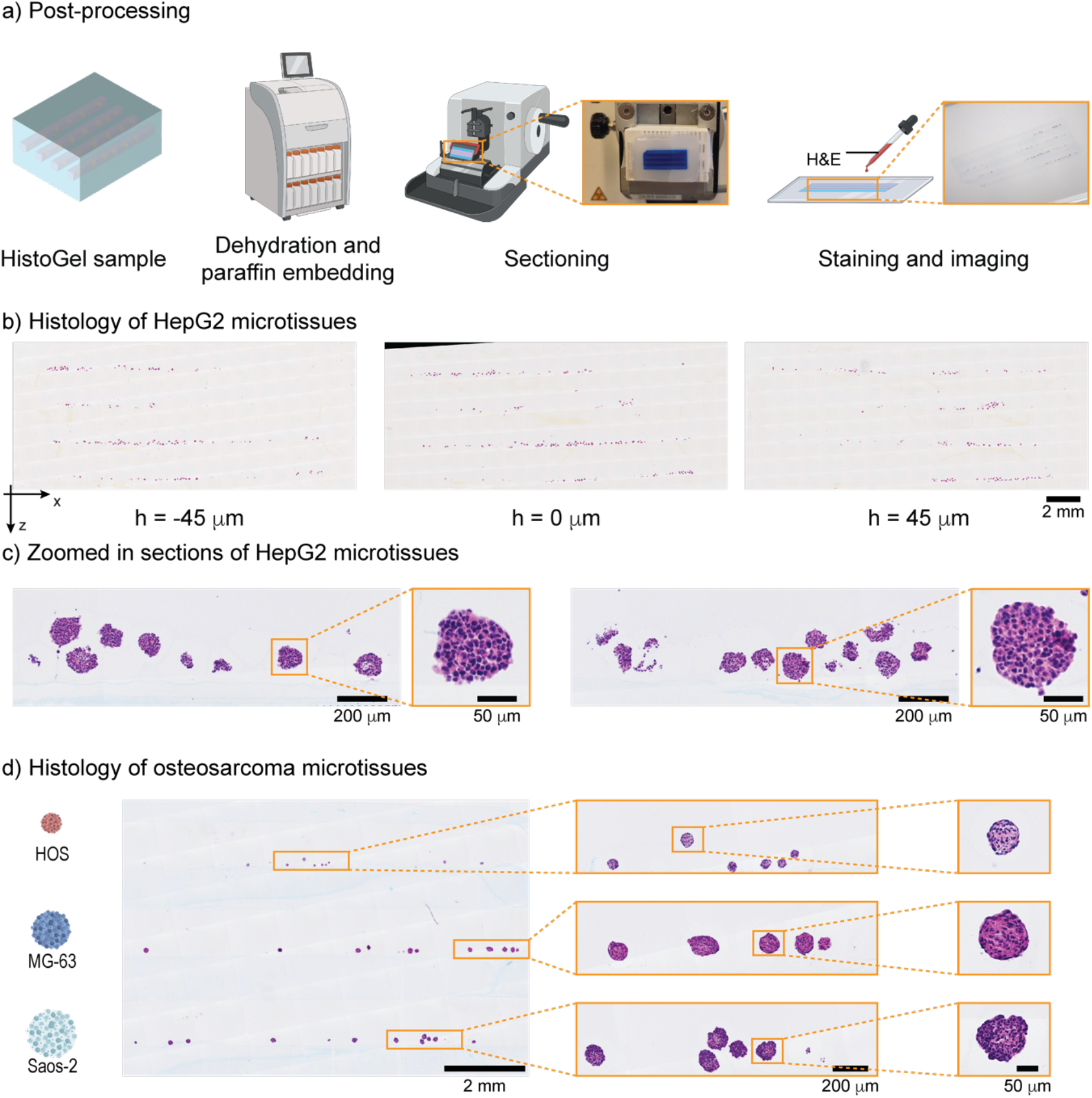
Acoustofluidic patterning enhances microtissue histology. **a)** Schematic of the histology process for paraffin infiltrated samples. The process consisted of sample dehydration by incubating in a series of ethanol and xylene solutions followed by paraffin embedding and sectioning with a microtome. Finally, the sections were stained and imaged. **b)** Microtissue histology (H&E staining) of acoustofluidically patterned and multiplexed, HepG2 microtissues within paraffin-embedded sections. The images correspond to serial sections spaced 45 μm apart. **c)** Zoomed-in-view of the section in b, to show the morphology of HepG2 microtissues. **d)** Paraffin sections containing three types of osteosarcoma cell lines (HOS, MG63, and Saos-2) stained with H&E. Patterned samples with each of these microtissues were multiplexed within one HistoGel slab (left) Zoomed in view of each spheroid type (center and right).

Our process for improved microtissue histology was designed to be compatible with standard histology workflows with minor adjustments. The dehydration protocol was modified to minimize shrinkage and maintain sample shape and aspect ratio. The constructs were the embedded in paraffin, sectioned, and finally, the sections were stained with H&E dyes and imaged (**Figure 4a**). The adjustments to the pre-processing of the constructs, like re-embedding into thicker HistoGel, did not affect their sectioning (**Figure 4a,b**). Further the acoustically patterned samples did not delaminate from the surrounding HistoGel.

Given that the construct positioned the microtissues within the multiplexed construct in the same plane, sections of samples containing each of four HepG2 spheroid samples were collected easily in the same sections (**Figure 4b**). Each section contained a high density of microtissues, demonstrating the utility of aligning the centers of mass of the microtissues to the same plane (same position in the y-axis) to improve microtissue histology. Within a given channel, microtissues were distributed across the z-axis and x-axis given the stochastic nature of sample loading in the acoustofluidic channel and that the pressure node was only driven to the preferred point in the y-axis. The coplanar alignment of the centers of mass enabled facile analysis of multiple samples within each section and the distribution across the z- and x-axes enabled visualization of individual microtissues (**Figure 4c**). Further, the microtissues remained intact morphologically in the final sections (**Figure 4c**).

As designed, the acoustofluidic patterning successfully generated multiplexed constructs with microtissues from distinct samples in the same, pre-defined plane of the final construct. We further analyzed the slices adjacent to the pre-defined central plane to assess minor variations in the location of the HepG2 microtissues in the final construct (**Figure S9**). Variations in the microtissue density from left to right along the slice indicate a tilt in the section plane, which was likely caused by a misalignment of the blade with the paraffin sample during sectioning. To correct for this, we considered microtissue height distributions within subsections of 2.5 mm x 3 mm and demonstrated that all microtissues were present within 161.2 ± 46.6 µm for these subsections (**Figure S9b**). Overall, this confirmed the robust patterning of the centers of mass of all microtissues to a narrow region of the final construct, consistent with the patterning efficiency observed after acoustic manipulation. Further, the minor deviations introduced by misalignment still allowed all microtissues to be sectioned and imaged within a narrow range in the y-axis of the final embedded construct, dramatically increasing the efficiency of the sectioning and imaging.

To demonstrate the utility of our method to pattern, process, and analyze multiplexed samples of more delicate microtissues, we applied the acoustofluidic patterning to microtissues made of three distinct osteosarcoma cell lines (**Figure 4d**). The HOS, MG-63, and Saos-2 microtissues form with different sizes and morphologies, emphasizing the need for mass-centric patterning across a multiplexed sample in order to image each of the samples within a given section. The microtissues were patterned and embedded as described above, and sections taken at the central plane of the final construct contained microtissues from each of the three samples. Further, all of the osteosarcoma microtissues remained intact during the patterning, embedding, and processing, allowing for comparative analysis.

## Conclusion

Analyzing microtissues throughout their lifecycle is essential for extracting high-content biological information, but their small size, high cell density, and random positioning within embedding media pose significant challenges for histological analysis. Current methods often require extensive sectioning and imaging, increasing time, cost, and complexity, while traceability across sections remains difficult. Our method addresses these challenges by enabling the precise, coplanar arrangement of microtissues within HistoGel. Approximately 80% of microtissues were positioned within 100 μm of the central plane of the sample. The method was validated using liver and osteosarcoma microtissues of varying sizes and compositions, demonstrating its robustness and versatility. Importantly, this technique integrates seamlessly into standard histological workflows, enabling improved sample preparation without altering downstream processing. While currently manual, this method holds significant potential for automation to further enhance efficiency and scalability. Future efforts, including numerical simulations of acoustic forces acting on microtissues, will refine device design and expand its applications. This acoustofluidic approach offers a transformative solution for microtissue histology, paving the way for more efficient, scalable, and high-throughput analysis of 3D in vitro models.

## Materials and methods

### Acoustofluidic device fabrication

We machined the acoustofluidic device by milling a cavity into an Aluminum plate (40 x 70 x 3 mm). A rectangular cavity was achieved by milling the aluminum plate (width = 1.1 mm, length = 40 mm, and depth = 1.5 mm) (**Figure S6**). For the initial iterations of the device, a depth of 1.1 mm was milled to produce a square cavity. We attached the piezoelectric transducer (PZ26, 3 mm x 20 mm x 40 mm, MEGGITT, Denmark) to the acoustofluidic device on the side using a cyanoacrylate-based glue (PST2C, Pattex Sekundenkleber Classic Matic, Germany). An insulated copper wire (l = 20 cm) was used to connect the transducer electrodes to the signal generator (AFG 3022B; Tektronix, USA) via the power amplifier (2100L-1911; MKS Instruments, Germany). To avoid reaching the Curie temperature, we used conductive silver (LS200N BC, Hans Wolbring GmbH, Germany) to connect the wires to the transducer instead of soldering. All the signals were monitored on an oscilloscope (LeCroy WaveSurfer 424, China)

### Acoustic device characterization

To find the resonant frequencies of the devices, baking yeast (fresh baking yeast, Migros, Switzerland) was used as tracker particles to perform a frequency sweep from 500 kHz to 900 kHz at 20 Vpp to search for the first acoustic mode. A sine wave with frequency modulation (Vpp = 20 V, Frequency Modulation: 20 kHz deviation at the rate of 100 Hz) was produced using a signal generator (AFG 3022B; Tektronix), coupled with an amplifier (NF Amplifier HSA 4101, Japan). Similar signals were used for patterning experiments of microtissue suspended in PBS (10010031, Thermofisher, USA). For experiments with HistoGel (22-110-678, Fisher Scientific, USA), a higher voltage was used to speed up the patterning (Vpp = 30 V, Frequency Modulation: 20 kHz deviation at the rate of 100 Hz).

### Numerical simulations using COMSOL Multiphysics

To demonstrate the rationale behind the geometry of the device, we performed a numerical simulation using COMSOL multiphysics 5.6. The cross section of the device was modeled in 2D using dimensions of the device. We performed a frequency domain study using Pressure Acoustics (Water/HistoGel), Solid Mechanics (Aluminum, Glass, and piezoelectric transducer), and Electrostatics (piezoelectric transducer). Acoustic-structure boundary was defined at the interface of the fluid domain and glass or aluminum domains and Piezoelectric effect was used for the transducer. Two different fluidic cavities were modeled. A square domain (1.1 mm x 1.1 mm) and another rectangular domain, similar to the device design (1.1 mm x 1.5 mm). The corners of the fluidic cavities were rounded off using a filet of 10 μm to avoid singularities at the corners. Frequency domain study was performed between 630 kHz and 670 kHz with a spacing of 1 kHz. Standard material properties were used from the COMSOL, except piezoelectric transducers (obtained from the manufacturer MEGGITT, Denmark). Frequencies corresponding to resonance frequencies in both cases were plotted.

### Temperature control

The temperature control system for the experimental setup consisted of a resistive heater (HT24S, Thorlabs) and a Pt100 temperature sensor (Distrelec, Switzerland). The system used a PID controller to maintain the desired temperature with high precision. The controller was interfaced with an Arduino Uno microcontroller to adjust the heater based on real-time feedback from the temperature sensor. The key parameters for the PID controller were determined using the Ziegler-Nichols method, optimizing the proportional, integral, and derivative coefficients to ensure stable and responsive control. The heater was in contact with the acoustofluidic device, maintaining a constant temperature of 45 °C, which is sufficient to prevent gelling of HistoGel during the patterning process. The temperature of the system was continuously monitored and logged using a computer interface.

### Microtissue patterning in HistoGel

The temperature control setup, consisting of a heater and temperature sensor was attached on the acoustofluidic device to keep the HistoGel (22-110-678, Fisher Scientific, USA) liquid while applying the acoustic signals. We filled the acoustic cavity with microtissues suspended in HistoGel, while the

λ/2 mode was active. 65 wt% HistoGel (in PBS) was used for the experiments as it was found to be suitable for acoustofluidic patterning and to hold shape during the post-processing steps. Once the cavity was filled, a glass slide was placed over the channel and clamped using paper clips. Microtissues were patterned first with λ/2 mode (f = 650 kHz) and then 3λ/2 (f = 2150 kHz) mode after ∼ 30 seconds to improve the patterning.

To solidify the HistoGel, the device was allowed to cool with the heater turned off. To speed up the cooling process, a cold aluminum block (dimensions: 40 x 20 x 60 mm; cooled to -20 °C) was placed over the device, while monitoring the temperature throughout the experiment. Once solidified, the samples were carefully retrieved from the channel using a spatula.

For post-patterning embedding, the samples were rotated 90° and placed inside custom-built aluminum molds, designed to accommodate four different samples within the same HistoGel slab (**Figure 3d**). This arrangement ensured that all samples were surrounded by HistoGel on all sides (**Figure S7**) and facilitated tracking of sample positions throughout post-processing. HistoGel (65 vol% in PBS) used for embedding was dyed with a blue tissue dye (YBP-1163-5, Bradley Products, USA) to distinguish it from the patterned samples, which allowed for easier sectioning and more accurate identification during imaging

### Patterning efficiency

The patterning efficiency of the device was evaluated after cooling down the HistoGel sample with microtissues patterned within. A video of the entire device was taken for each of the samples. From the videos, a series of overlapping images were obtained for the entire channel. The images were merged together in Adobe Photoshop using the photomerge option (**Figure S8a**). Parts of the image that could not be merged using photoshop were merged manually. The channels were rotated by using the straighten feature of the crop tool in Photoshop. Images were then imported into Fiji, where the center of each microtissue was marked manually and their positions were stored (**Figure S8b**). The positions of the microtissues were analyzed and microtissues were defined as successfully patterned if their center lay within 100 μm of the center of the channel.

### HistoGel viscosity measurement

To measure the viscosity, we used shear rheology on 65% HistoGel at different temperatures (75, 65, 55, 45 °C). A shear rate ramp (from 0.1 s^-1^ to 1000 s^-1^) was performed at each temperature using parallel plate geometry (Ø = 20 mm; PP20, Anton Paar) with 0.5 mm gap. The measured values where torque was less than 100 nN were ignored. Zero-shear viscosity values were estimated as an average of the viscosity values between 10 s^-1^ to 100 s^-1^, as the plot shows a plateau here.

### Speed of sound measurement setup

A 5 MHz longitudinal transducer/receiver (Panametrics V109, USA) was connected to an ultrasound pulse receiver (Panametrics Model 5800, USA). The transducer was placed over the sample using a custom made PMMA well with known dimensions (10 mm), with a steel plate at the bottom. An oscilloscope (LeCroy WaveSurfer 424, China) was used to acquire the signals. A pulse emitted from the transducer passes through the liquid and is reflected by the steel plate. Measuring the difference between the emitted pulse and the received pulse gives the speed of sound in the fluid. The following parameters were used to acquire the measurements, PRF = 125 Hz, energy = 50 μJ, damping = 15 Ohm, HP filter = 300 kHz, LP filter = 20 MHz, input and output attenuation = 0, Gain = 20 dB, sensitivity = 65.9 dB. To maintain the temperature of the measurement setup we attached the heater and temperature sensor on the steel plate.

### HepG2 microtissue formation and fixation

Human hepatocellular carcinoma (HepG2) (HB-8065™, ATCC, USA) cells were cultured in Dulbecco’s Modified Eagle Medium (DMEM; 10313021, Thermofisher, USA) supplemented with 10 % fetal bovine serum (FBS; A5670401, Thermofisher, USA) and 1 % penicillin-streptomycin (P/S; 15070063, Thermofisher, USA). Cells were cultured in 75 cm2 flask and were passaged when 80% confluency was reached. HepG2 microtissues were cultured in Sphericalplate 5D platform technology (SP5D, Kugelmeiers Ltd., Switzerland) with a starting cell density of 200 cells per micro-well. After six days, the microtissues reached an average diameter of 150 μm and were harvested and fixed with 4 % paraformaldehyde (PFA) for 20 minutes at room temperature. The fixed microtissue were stored in phosphate-buffered saline (PBS) at 4°C.

For the formation of HepG2 microtissues with a diameter of 250 μm and 500 μm, the cells were seeded with respective density of 200 cell/well and 500 cell/well in a 96-well Ultra-Low Attachment Surface plate (ULA; 7007, Corning, USA). After five days, the microtissues were harvested and fixed with 4 % paraformaldehyde (PFA) for 20 minutes at room temperature. The fixed microtissue were stored in phosphate-buffered saline (PBS) at 4°C.

### Osteosarcoma microtissue formation and fixation

Human osteosarcoma HOS (CRL1543, ATCC, USA), MG63 (Mayo Clinic, USA) and Saos-2 cells (ACC 243, Leibniz Institute DSMZ, Germany) were cultured DMEM-F12 Ham mixture (D8437, Sigma-Aldrich, USA), supplemented with 10% FBS (F4135, Sigma-Aldrich, USA), 2mM L-Glutamine (G7513, Sigma-Aldrich, USA), and 1% P/S (P4333, Sigma-Aldrich, USA). They were cultured in T-flasks at 37°C in a humidified atmosphere with 5% CO2. The culture medium was replaced every 3 days, and the cells were subcultured when they reached 70-80% confluency. Cells were detached from T-flasks and suspended in microtissue medium made of DMEM/Ham’s F-12, 10% horse serum (HS; P30-0702, Pan Biotech GmbH, Germany), 25mM HEPES buffer (15630-056, Gibco, USA), and 1% P/S (P4333, Sigma-Aldrich, USA) seeded at the following densities: HOS, 2.5 x 10^4^ cells/ml; MG63, 2.5 x 10^3^; SAOS-2, 2 x 10^4^ cells/ml. 100 μl of cell suspension was dispensed into each well of round U-bottom ULA 96-well plates. Microtissues were cultivated for up to 7 days with a medium exchange at day 4. After 7 days of culture, the microtissues reached a diameter of 122.7 ± 9.2 um for HOS, 291.9 ± 27.8 um for MG63, and 258.9 ± 15.9 um for SAOS-2. The microtissues were harvested on day 7 and fixed with 4 % PFA for 1 hour at room temperature. The fixed microtissues were then stored in HBSS (+/+; Gibco, USA) at 4°C until use.

### Paraffin embedding and sectioning

The HistoGel samples were embedded in paraffin by first dehydrating in solutions of ethanol and xylene before being infiltrated with paraffin as per the protocol in **Table 1**. Finally, the samples were embedded in paraffin using standard laboratory procedures. Sections of 5 um were cut with a Leica RM2265 microtome for sectioning. The samples were first trimmed until the undyed channel appeared and 300 µm more were removed. Afterwards, the appearance of microtissues was monitored with a microscope at intervals of every 15 µm (i.e. 1 section out of 3 was checked). When the first microtissues were spotted, sections every 15 µm (1 section over 3) were collected.

**Table 1.** Protocol used for paraffin embedding.

| Solution | Time (min) |
| --- | --- |
| Ethanol 70% | 45 |
| Ethanol 80% | 45 |
| Ethanol 95% | 90 |
| Ethanol 100% | 45 |
| Ethanol 100% | 45 |
| Ethanol 100% | 45 |
| Xylene | 45 |
| Xylene | 60 |
| Xylene | 75 |
| Paraffin | 75 |
| Paraffin | 135 |

### Hematoxylin and Eosin staining

To localize the microtissues, slides were stained with hematoxylin and eosin, H&E. The slides were equilibrated and dried at room temperature for 1 hour, then rehydrated in deionized water for 10 min. The slides were then incubated in Harris Hematoxylin (3873.2500, Biosystems) for 5 min and washed with running tap water. They were then differentiated for a few seconds in 1 % acid-alcohol (Absolute alcohol: 7:10, VWR 20820.362, Hydrochloric acid 37%: 0.1:10, Sigma Aldrich, 30721 and H2O MiliQ 2.9:10) and washed under running tap water for 10 min. The slides were incubated in Eosine-Phloxine solution for 1 min (Eosin: 1:100, E4382, Sigma Aldrich, USA; Phloxine: 1:100, P2759, Sigma Aldrich, USA) and washed the last time. Finally, the slides were mounted with Eukitt (03989, Sigma Aldrich, USA) and dried overnight at room temperature

### Image acquisition

Images of the H&E-stained sections were acquired with an Olympus VS200 slide scanner. An overview image of the entire glass slide was acquired at 10x (UplaXapo, NA=0.40) magnification in brightfield.

### Histological sections image analysis

Python libraries were used to process the images of the histological sections.The goal is to identify on which sections the microtissues start to appear and disappear in function of their location on the section. First, using edge detection, the images are cropped and rotated to align the sections vertically. Second, the sections were divided in 40 regions of 2.5 x 3 mm^2^ (**Figure S9a**). Third, the image was segmented using a binary mask based on the distinctive pink color of the microtissues. Finally, the aligned binary images are saved with the grid visible. On the processed image, the microtissues appear white while the background is in black. The code used to process the images is available in the supporting information.

For each grid position, the depth of the first section (S_first_) and the last section (S_final_) on which microtissues appear were manually found. The interval thickness containing microtissues is calculated as followed: (S_final_ - S_first_ + 1)*15. The section interval is multiplied by 15 µm as 1 section is collected every 15 µm. The depth of the interval center is calculated as followed: [S_first_+(S_final_ - S_first_)/2]*15.

## Contributions

D.V.D.: Methodology, experiment development, acoustofluidics experiments, data compilation, analysis, writing manuscript. E.V.d.B.: Microtissue formation, histology experiment development and analysis, writing – manuscript. S.J.: Experiments, writing manuscript. N.G.: Device design development and testing, patterning experiment analysis in water, embedding mold formulation and testing. C.F.: Early device iteration development and testing, heater integration. A.E.: Early device iteration testing, patterning efficiency algorithm formulation. E.B.: Microtissue formation and analysis, writing – manuscript. M.R.: Microtissue formation and analysis, writing manuscript. S.H.: Project formulation, histology experiment support. D.M.: Initial idea formulation, early sectioning trials, data analysis. S.B.P.: Planning, writing manuscript. G.W.: Project formulation, microtissue formation and histology experiment support. J.D.: Project formulation, acoustofluidics experiment support, device fabrication, writing manuscript. M.W.T.: Project formulation, device fabrication support, device design formulation, writing manuscript.

## Acknowledgements

This work was supported by a BRIDGE Discovery Grant (40B2-0_211752). Aspects of Figure 4 were made with BioRender.com.

## Notes

### Competing Interest Statement

The authors have declared no competing interest.

