## Supporting Data for "Acoustofluidic patterning for improved microtissue histology"

### Supporting Information

**Table of Contents**

**Supporting Figure S1** **Alignment of particles in acoustofluidic channels……………………….……3**

**Supporting Figure S2** **Patterning of microtissues in the acoustofluidic cavity with and without glass cover…………………………………………………………………….4**

**Supporting Figure S3** **Temperature control in the acoustofluidic device………………………….5**

**Supporting Figure S4** **Viscosity of 65% HistoGel…………………………………………………...6**

**Supporting Figure S5** **Speed of sound in HistoGel…………………………………………………..7**

**Supporting Figure S6** **Acoustofluidic device design………………………………………………...8**

**Supporting Figure S7** **Sample re-embedding molds………………………………………………...9**

**Supporting Figure S8** **Calculating patterning efficiency…………………………………………..10**

**Supporting Figure S9** **Analysis of microtissue patterning in sectioned samples…………………11**

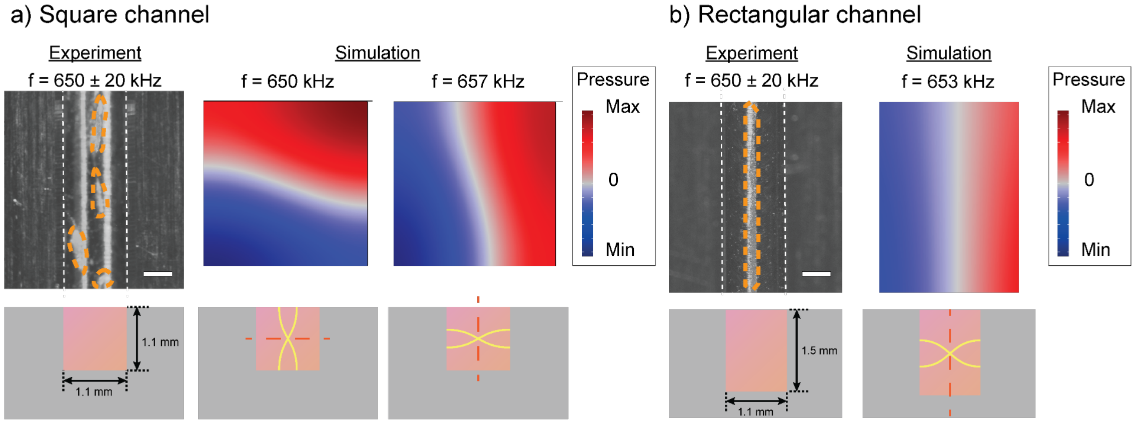


**Figure S1. Alignment of particles in acoustofluidic channels. a)** Square channel, particles (marked by orange) were either patterned at the center (y-axis) or levitated in the channel (z-axis). This was due to their $\lambda/2$ modes occurring close to each other. **b)** Rectangular channel, particles (marked by orange) were patterned at the center (y-axis). The $\lambda/2$ mode for levitation (z-axis) and patterning (y-axis) occur far apart from each other. The walls of the channel are marked with white dotted lines marked in **(a, b).**

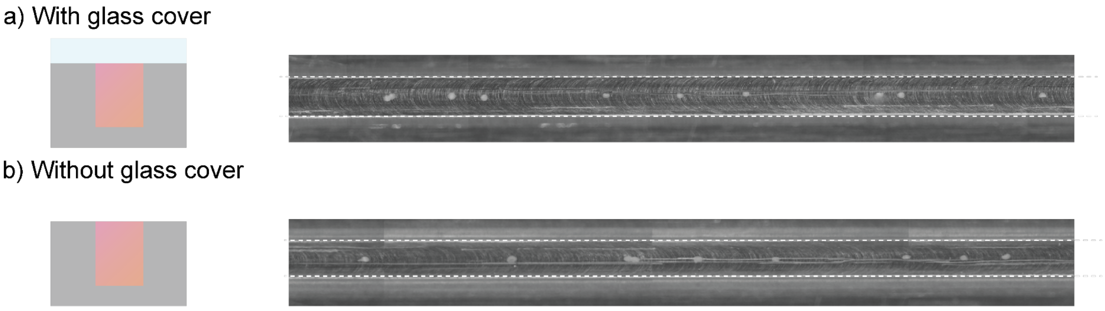


**Figure S2. Patterning of microtissues in the acoustofluidic cavity with and without glass cover. a)** Patterning without a glass cover. **b)** Patterning with a glass cover. The walls of the channel are marked with white dotted lines marked in **(a, b)**. While patterning was observed in both cases, the patterning efficiency (defined by the percentage of microtissues within 100 $\mu$m from the center of the channel) was higher without the glass cover. Since in the latter case, the acoustic field is turned on before the microtissue enters the acoustic cavity, they can avoid sticking to the cavity walls or the glass slide. For the subsequent experiments, we combined the two methods by turning on the acoustic signals while microtissues were filled into the acoustic cavity and later closing the cavity with the glass slide.

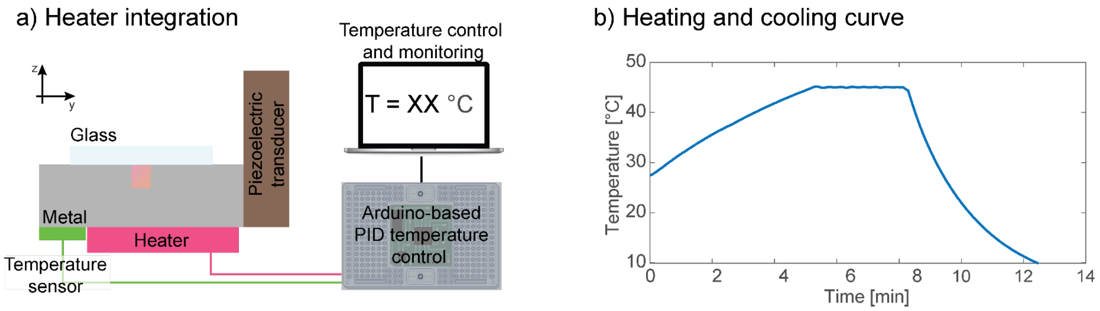


**Figure S3. Temperature control in the acoustofluidic device. a)** A heater and temperature sensor were integrated into the acoustofluidic device, they were controlled with a PID controller interfaced to a computer using Arduino. **b)** Heating and cooling curve during a typical experiment. Cooling was done by placing a cold metal block on the device.

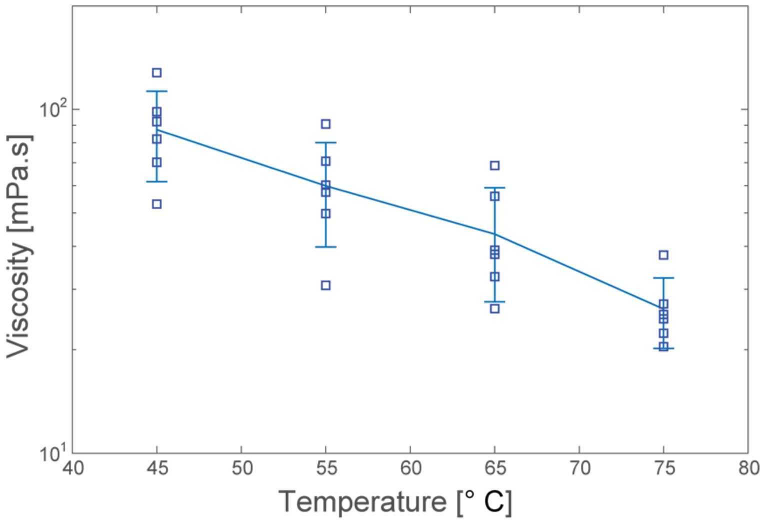


**Figure S4. Viscosity of 65% HistoGel.** Viscosity of HistoGel decreased with increasing temperature.

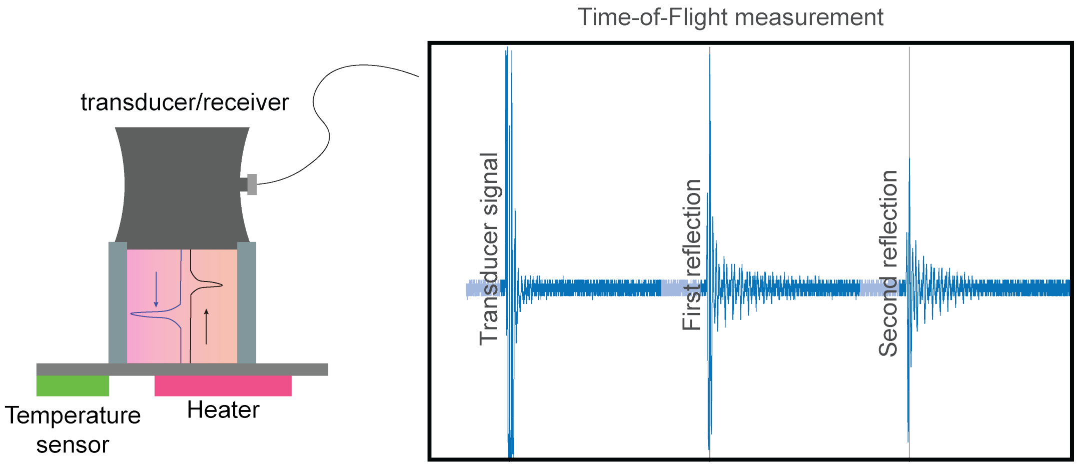


**Figure S5**. **Speed of sound in HistoGel.** Measurement of the speed of sound in HistoGel was done with an ultrasonic transducer and receiver to measure the time-of-flight.

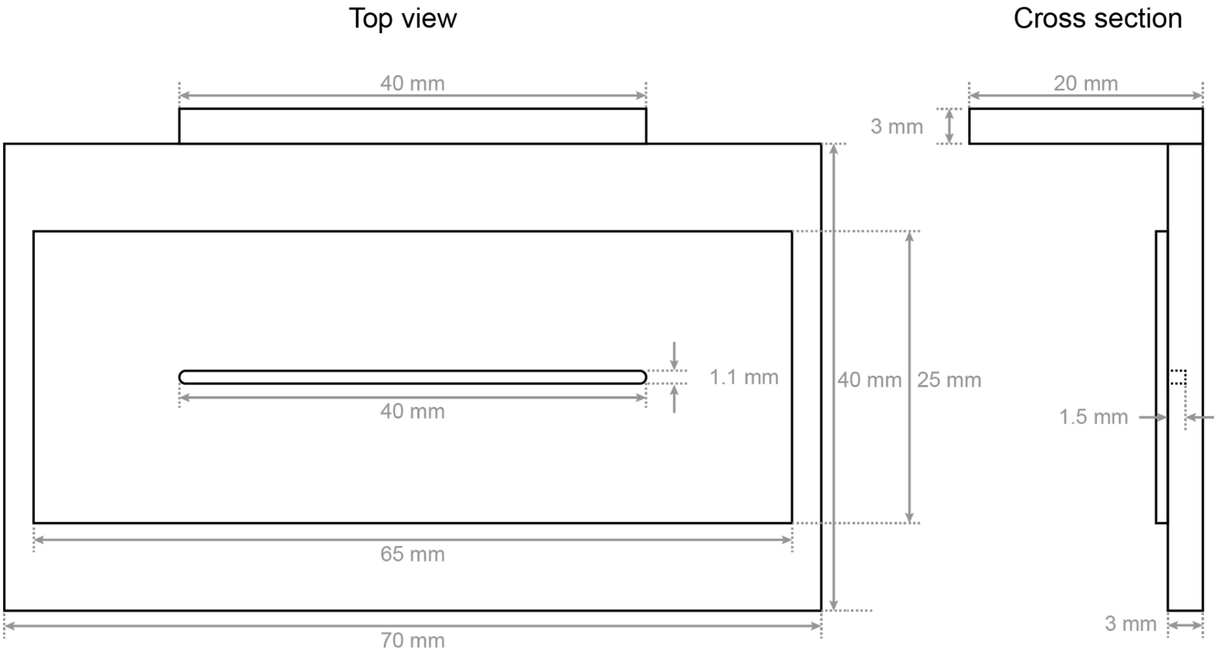

**Figure S6. Acoustofluidic device design**

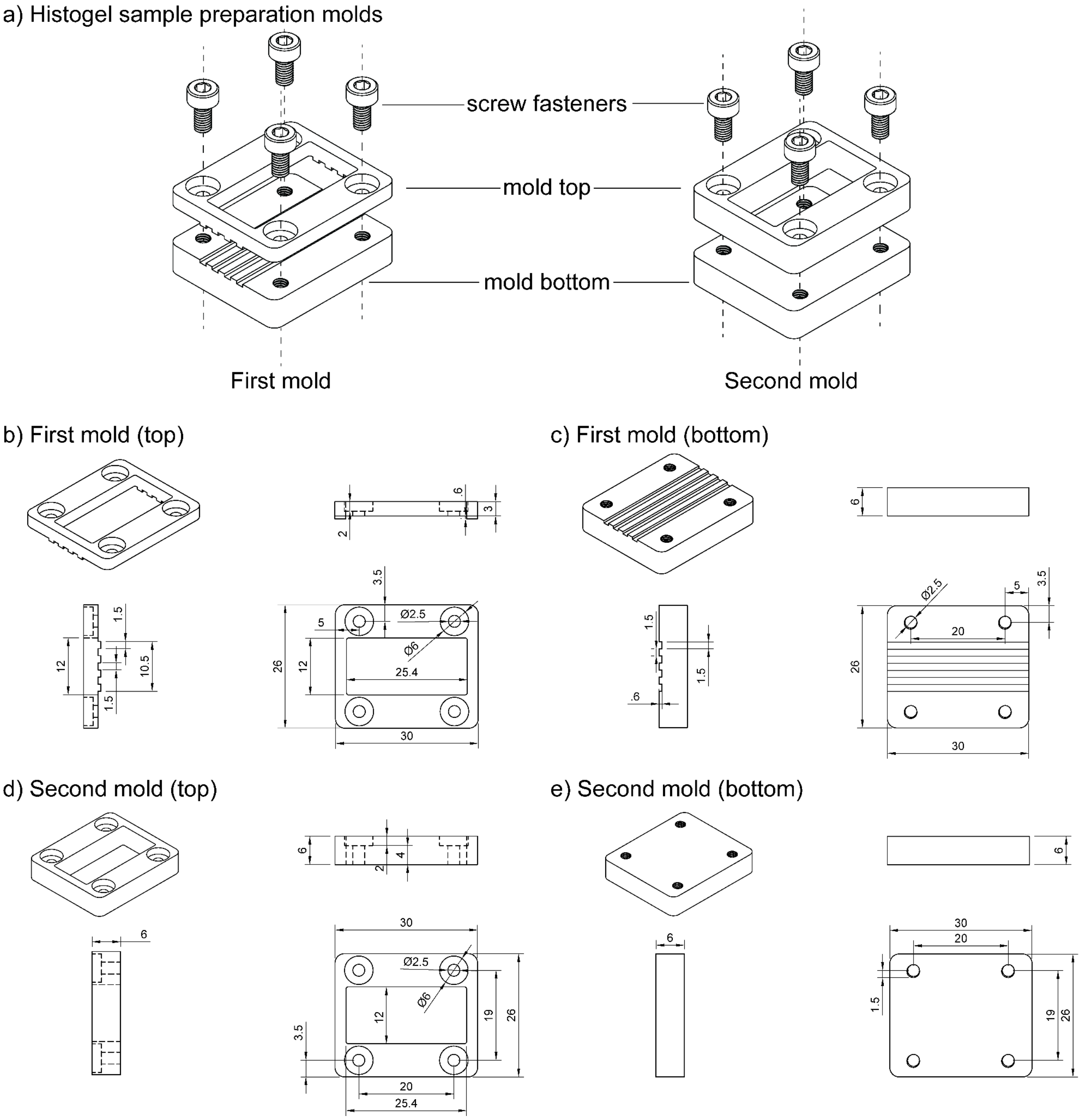


**Figure S7. Sample re-embedding molds**

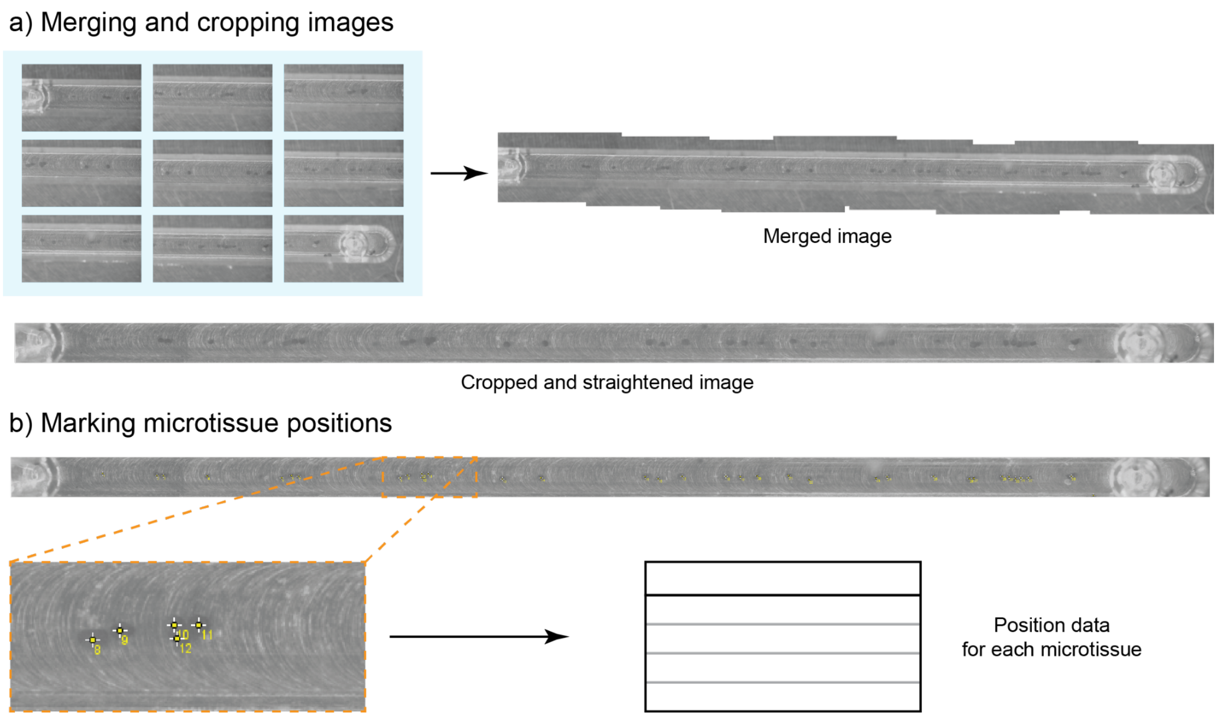


**Figure S8. Calculating patterning efficiency. a)** Images of the entire channel were merged together to form a continuous image of the channel. This image was straightened and cropped to the size of the channel. **b)** Every microtissue in the channel was marked and its positions were stored.

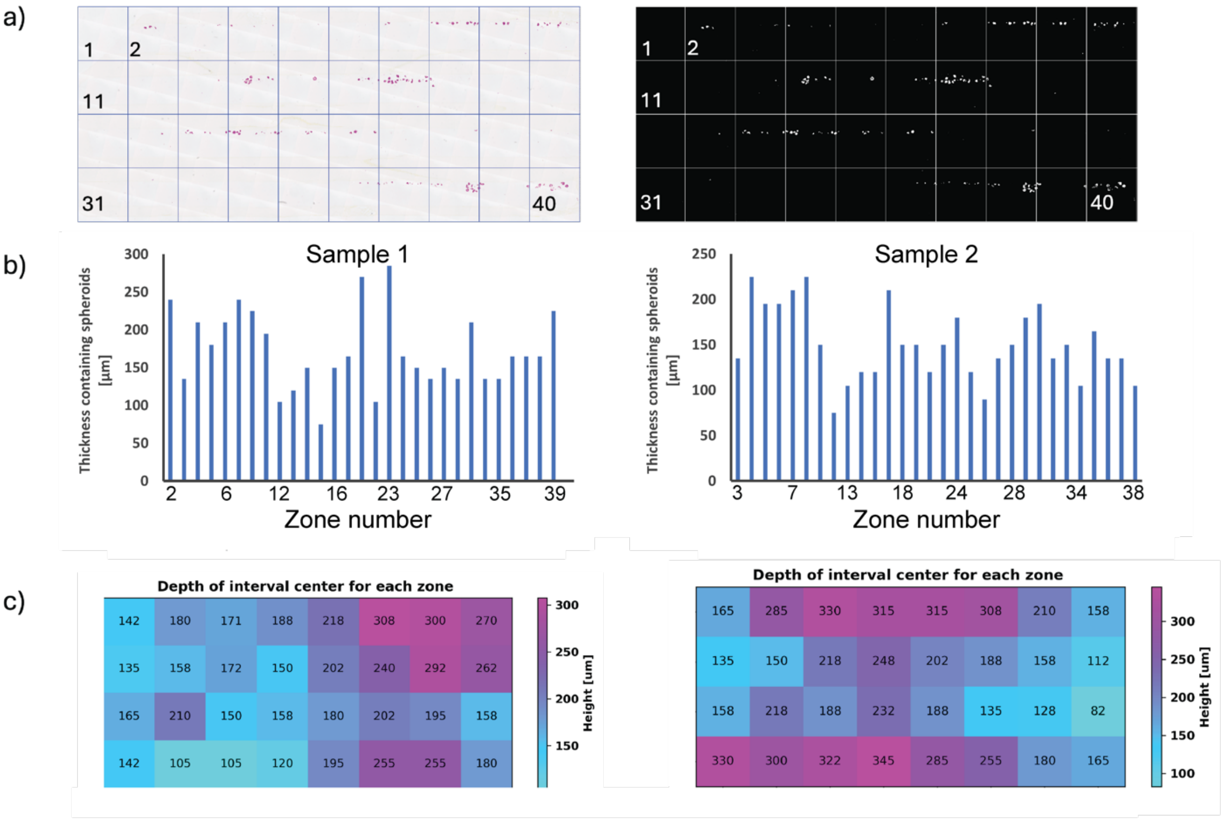


**Figure S9. Analysis of microtissue patterning in sectioned samples. a)** Histological section stained with H&E divided in 40 zones (left) and after segmentation (right). **b)** Thickness of the sample containing the spheroids for each zone for sample 1 (left) and sample 2 (right). **c)** Heat map showing the height of the center for each thickness containing spheroids for sample 1 (left) and sample 2 (right).
